# The utility of explainable A.I. for MRI analysis: Relating model predictions to neuroimaging features of the aging brain

**DOI:** 10.1101/2024.09.27.615357

**Authors:** Simon M. Hofmann, Ole Goltermann, Nico Scherf, Klaus-Robert Müller, Markus Löffler, Arno Villringer, Michael Gaebler, A. Veronica Witte, Frauke Beyer

**Author notes:** equal contribution (Simon M. Hofmann, Ole Goltermann). Corresponding authors: Simon Hofmann,; Ole Goltermann.

## Abstract

**Introduction:** Deep learning models highly accurately predict brain-age from MRI but their explanatory capacity is limited. Explainable A.I. (XAI) methods can identify relevant voxels contributing to model estimates, yet, they do not reveal which biological features these voxels represent. In this study, we closed this gap by relating voxel-based contributions to brain-age estimates, extracted with XAI, to human-interpretable structural features of the aging brain.

**Methods:** To this end, we associated participant-level XAI-based relevance maps extracted from two ensembles of 3D-convolutional neural networks (3D-CNN) that were trained on T1-weighted and fluid attenuated inversion recovery images of 2016 participants (age range 18-82 years), respectively, with regional cortical and subcortical gray matter volume and thickness, perivascular spaces (PVS) and water diffusion-based fractional anisotropy of main white matter tracts.

**Results:** We found that all neuroimaging markers of brain aging, except for PVS, were highly correlated with the XAI-based relevance maps. Overall, the strongest correlation was found between ventricular volume and relevance (*r* = 0.69), and by feature, temporal-parietal cortical thickness and volume, cerebellar gray matter volume and frontal-occipital white matter tracts showed the strongest correlations with XAI-based relevance.

**Conclusion:** Our ensembles of 3D-CNNs took into account a plethora of known aging processes in the brain to perform age prediction. Some age-associated features like PVS were not consistently considered by the models, and the cerebellum was more important than expected. Taken together, we highlight the ability of end-to-end deep learning models combined with XAI to reveal biologically relevant, multi-feature relationships in the brain.

## 1. Introduction

Deep neural networks (DNNs) have become a valuable asset for the analysis of neuroimaging data, facilitating the classification of brain-related diseases (e.g., Alzheimer or multiple sclerosis; Vieira et al., 2017; Zhang et al., 2018) and the estimation of continuous variables such as brain-age (BA; Ball et al., 2021; Bashyam et al., 2020; Cole et al., 2017; Dinsdale et al., 2021; Feng et al., 2020; Peng et al., 2021) from minimally processed magnetic resonance images (MRIs), avoiding a priori feature selections. While the models have shown to be highly accurate prediction models in many domains, their decision process is difficult to interpret due to the high complexity of these models (also known as the black box problem). Only recently the imaging community started to employ algorithms that partially explain the internal decision process of these models (Böhle et al., 2019; Eitel et al., 2019; Hofmann et al., 2022; Levakov et al., 2020; Thomas et al., 2019). The majority of these explainable A.I. (XAI) algorithms generate explanations *post hoc* (Ras et al., 2022) in the form of relevance maps (also heatmaps, or saliency maps) highlighting pixels or voxels which have been relevant for the prediction of the model. These relevance-based explanations became popular since they appeal to human intuition. Common benchmarks for both prediction models and XAI algorithms often contain natural images of, e.g., cats, cars and other familiar objects. When a prediction model correctly classifies samples from such familiar categories, the corresponding relevance maps highlight particular object-related features that usually appeal to our understanding of the corresponding class (e.g., the eyes and ears of a cat). However, in contrast to the application of XAI on natural images, our intuition regarding the relevance maps of MRI-based predictions is limited, even for neuroimaging experts. Moreover, there is a danger of human bias when selectively interpreting relevance maps (confirmation bias; Adebayo et al., 2020; Bacon, 1620; Ghassemi et al., 2021; Lakens, 2022). Therefore, there is a need for analyzing relevance maps quantitatively and relate them to human interpretable features. One way to achieve this is to aggregate relevance over regions derived from brain atlases. For instance, Eitel et al. (2019) used gray and white matter (WM) atlases to determine which brain regions were most relevant for their deep learning model to accurately diagnose multiple sclerosis. Another way is to overlap explanation maps with spatial maps of other neuroimaging markers, which also capture distributed neural-structural properties. This way, in our previous work (Hofmann et al., 2022), we could show that white matter hyperintensities (WMHs), a common marker of cerebral small vessel disease (cSVD; Cuadrado-Godia et al., 2018; Li et al., 2018) in WM, were relevant for our FLAIR-based BA prediction model as indicated by the XAI method Layer-wise Relevance Propagation (LRP; Lapuschkin et al., 2019). LRP takes a prediction and propagates it back through the DNN layer-by-layer, identifying the relevant contributions to the prediction up until the input (here the MRI). WMH only partially explained the highly accurate BA predictions of our model. That is, only a fraction of the total relevance scores were captured by voxels containing WMHs. Additionally, on visual inspection, we found that higher relevance reflected gray matter atrophy around the ventricles but did not quantify this yet. In summary, while previous studies demonstrated that XAI methods can provide initial information about the decision process of deep-learning-based BA models (Hofmann et al., 2022; Levakov et al., 2020), we so far only partially understand which neural features the highlighted areas represent.

In this study, we investigated the contribution of well-known quantifiable imaging markers of age-related biological changes to highly precise deep learning-based BA estimates. To this end, we tested gray matter (GM) properties, including GM volume (GMV) in the cortex, in subcortical regions, the cerebellum and brainstem, cortical thickness (CT), and ventricular volume as imaging markers of GM atrophy, a prominent feature of brain aging (Fjell & Walhovd, 2010). Moreover, we examined the WM microstructure of neural fiber bundles represented in fractional anisotropy measures (FA; Basser & Pierpaoli, 2011), which is known to reflect age-related changes in the WM organization (Schilling et al., 2022). Finally, we investigated another imaging marker of cSVD, namely perivascular spaces (PVS), which have a characteristic appearance of small, liquid-filled structures parallel to perforating blood vessels (Swieten et al., 1991; Wardlaw et al., 2020). The automatic quantification of PVS is relatively novel, but it has been shown that age is a strong predictor for the presence of PVS (Francis et al., 2019).

Based on these associations, we hypothesized that higher relevance scores (i.e., higher predicted BA) are associated with smaller CT in cortical, and lower GMV of both cortical and subcortical regions, as well as of cerebellum and brainstem. Moreover, we expected higher relevance values to overlap with PVS in deep WM around the basal ganglia and in the whole brain and to correlate with lower FA values in WM tracks.

## 2. Methods

### 2.1 Participants

Participants took part in the baseline assessment of the population-based cohort LIFE-Adult study (Loeffler et al., 2015), in which they underwent extensive clinical screenings, including measures of height, weight, blood pressure, blood-based biomarkers, neuroimaging, cognitive performance and surveys on mental health and lifestyle (for further information, refer to Loeffler et al., 2015). Among the more than 10,000 subjects enrolled in the LIFE Adult study, 2637 participants underwent a baseline 1-hour brain MRI recording session. Out of those individuals who received MR-scans, 621 participants were excluded primarily due to pathologies, resulting in 2016 subjects (age range of 18–82 years, mean age = 57.32, median age = 63.0; n_females_ = 946) for further analysis. Among the 621 participants excluded, exclusion criteria partially overlapped: Exclusion criteria included previous strokes (n = 54), excessive brain lesions evaluated by trained medical professionals (n = 114), including white matter hyperintensities (WMH) with a Fazekas score of 3 (n = 44), radiological diagnosis of brain tumors (n = 22), diagnosis of multiple sclerosis (n = 5), epilepsy (n = 27), recent cancer treatment (n = 109), centrally active medication (n = 275), cognitive impairments indicated by an MMSE score < 26 (n = 80), and inadequate quality MRIs (failing a visual quality check, e.g., motion artifacts, n = 41).

### 2.2 MRI data acquisition

MRI data was obtained on a 3T Siemens Verio scanner, equipped with a 32-channel head coil. In this study, we made use of three MRI-sequences commonly utilized in clinical settings: i) Structural T1-weighted images were captured using a MPRAGE-sequence (T1w; 1 mm isotropic voxels, 176 slices, TR = 2300 ms, TE = 2.98 ms, TI = 900 ms, field of view 256×240×176 mm^3^). ii) Fluid-attenuated inversion recovery images (FLAIR; 1 x 0.49 x 0.49 mm voxels, sagittal orientation, 192 slices, TR = 5000 ms, TE = 395 ms, TI = 1800 ms, field of view 250×250×192 mm^3^). iii) Diffusion-weighted images (DWI; double spin-echo sequence, voxel size = 1.7 × 1.7 × 1.7 mm^3^, TR = 13.8 ms, TE = 100 ms, field of view 220×220×123 mm^3^, matrix = 128×128, maximum *b*-value = 1000 s/mm^2^ in 60 directions, and 7 volumes with *b*-value = 0 s/mm^2^).

### 2.3 MRI preprocessing

T1w images were processed with FreeSurfer v.5.3.0. We used cortical surface parcellations for relevance extraction and to determine metrics for both cortical thickness (CT) and cortical gray matter volume (GMV) according to the Desikan-Killiany atlas (DK; Klein & Tourville, 2012). Similarly, FreeSurfer’s automated subcortical segmentation procedure (Fischl et al., 2002) was employed to obtain subcortical GMV bilaterally (putamen, nucleus accumbens, hippocampus, amygdala, globus pallidus, caudate, thalamus) and 4 cerebrospinal fluid (CSF) volumes (lateral ventricle, inferior lateral ventricle, third ventricle and CSF), brainstem volume and WM and GM volume of the cerebellum. Freesurfer results were visually checked according to Klapwijk et al. (2019) which led to the exclusion of 24 individuals from cortical and subcortical analyses mostly based on segmentation errors due to high head motion. T1w imaging was also used to detect perivascular spaces (PVS) using the SHIVA-PVS toolbox (Boutinaud et al., 2021; https://github.com/pboutinaud/SHIVA_PVS; accessed in March 2024). This toolbox utilizes a deep learning-based algorithm designed for the detection and segmentation of PVS. It is constructed upon a 3D-convolutional neural network (CNN)-based autoencoder including a U-shaped network (U-Net; Ronneberger et al., 2015) trained on either T1w or both T1w and FLAIR co-registered images. We employed the multimodal T1-FLAIR model to segment PVS for each subject on the T1w image in the individual’s FreeSurfer space. The PVS segmentation failed in 4 individuals which were excluded for further analysis. The resulting PVS probability maps were thresholded at 0.5. In order to make sure we captured the relevance attributed to the border region of the PVS, we additionally created maps in which we dilated PVS by one and two voxels, respectively. We used whole-brain PVS segmentations as well as PVS masked around the basal ganglia (defined using the ATAG atlas, dilated by 5 voxels, https://www.nitrc.org/projects/atag), because of their larger size and more reliable detection (Boutinaud et al., 2021). DWI were processed using a customized pipeline including the in-house software LIPSIA, MRTRX v.3.0 and the FMRIB Software Library FSL, v.5.0.9 (Jenkinson et al., 2012; Lohmann et al., 2001; Tournier et al., 2019). Initially, the raw data for each participant underwent denoising and the reference (B0) images were processed with Gibbs unringing to improve data quality. Skull-stripping was performed using the FSL function BET and outliers were replaced using FSL eddy outlier replacement. Finally, motion correction and distortion correction was performed and the B0 image was coregistered to the T1w images within LIPSIA. After upsampling the data to 1 mm, diffusion tensor fitting was performed in LIPSIA. The resulting FA image was coregistered to the MNI space using ANTs (v.0.4.2; Avants et al., 2008) in Python v.3.10 and thresholded at ≥ 0.2 to exclude extreme cross-subject variability within small individual WM tracts and to mitigate misalignments during registration. We then parcellated the FA maps into 48 regions using the DTI-based WM atlas JHU (Mori et al., 2005) and calculated the mean FA for each region. FA images were not available in 17 individuals.

### 2.4 Explainable deep learning predictions of brain age

#### 2.4.1 Prediction models

Brain age (BA) prediction models were based on a multi-level ensemble (MLENS) architecture consisting of three sub-ensembles, each built on a specific MRI sequence (T1w, FLAIR, susceptibility weighted images; SWI). Each sub-ensemble comprised ten independently trained base models, each of which was constructed using a 3D-CNN implemented in Keras (v.2.3.1; Chollet, 2015). The base model architecture involved five convolutional blocks, followed by leaky rectified linear units (ReLUs) and max-pooling layers. The convolutional blocks were followed by dropout regularization and two fully connected layers, with the output layer bias set to the mean age in the dataset. The network was trained to minimize mean squared error (MSE) using the ADAM optimizer. The dataset was divided into training, validation, and test sets in an 8:1:1 ratio, with model performances evaluated on the test set and reported as mean absolute error (MAE) for better interpretability. Subsequently, a linear head model with weight regularization (*L^2^-norm*) was trained on the predictions of the 10 base models per sub-ensemble on the validation set, followed by evaluation on the test set. The resulting predictions from these three sub-ensembles on the test set were then utilized to train an additional head model atop the MLENS using a 5-fold cross-validation procedure, thereby obtaining aggregated predictions across all MRI sequences. A more detailed description of the model architecture and model performance can be found in Hofmann et al. (2022).

#### 2.4.2 Explanation algorithm and relevance maps

We applied layer-wise relevance propagation (LRP; Bach et al., 2015; Lapuschkin et al., 2019; Montavon et al., 2018; Samek et al., 2021) to study which features drive the performance of our BA prediction model. LRP is an algorithm that generates explanatory heat maps in the input space (i.e., relevance maps) of machine learning models, including non-linear deep learning models. This is achieved by decomposing the prediction *f(x)* of the model *f* with respect to the input *x* into relevance scores *R*. In deep learning models, this is computed layer by layer down to the input space, while satisfying the conservation criterion in each network layer: *ΣR* = *f*(*x*) (for details, see Montavon et al., 2019). Note, as a consequence of this property, the same prediction *f(x)* (here, the age estimate) for brains and their regions of different sizes, leads to different expected relevance scores per voxel. That is, the larger the brain (region) the more the total relevance (*ΣR*) is spread over voxels. Therefore, for statistical comparisons across participants, we used the sum-relevance over the expected (i.e., mean) relevance to account for brain size. Unlike alternative explanation methods, LRP stands out for its computational efficiency and its ability to integrate both local and global feature interactions, which are essential for accurate model prediction. In our study, we applied LRP on T1 and FLAIR sub-ensembles, obtaining relevance maps for all subjects and image modalities in the FreeSurfer subject space. Among the 2016 subjects, in 137 and 140 LRP was not performed for the T1 and FLAIR sub-ensembles due to computational reasons, respectively. These groups did not differ significantly in age (two-sample t-test: T_T1_ = -0.44, p_T1_ = 0.6; T_FLAIR_ = -0.9, p_FLAIR_ = 0.5) and sex (Chi-square_T1_ = 0.1, p_T1_ = 0.7; Chi-square_FLAIR_ = 0.05, p_FLAIR_ = 0.8) from the rest of the sample. Relevance maps were also transformed to MNI space using ANTs as described in Hofmann et al. (2022). As a result of LRP, each voxel is assigned with a relevance score where negative values represent model evidence in the input towards a younger age (below the mean age in the cohort), and positive relevance values represent evidence towards a higher age (above the mean age in the cohort).

### 2.5 Associations of brain age relevance maps with imaging features

To evaluate the biological relevance of our deep learning-based BA model, we examined different aging-related imaging markers derived from T1w, FLAIR, and DWI images (for an overview see **Fig.1**).

**Fig. 1:**
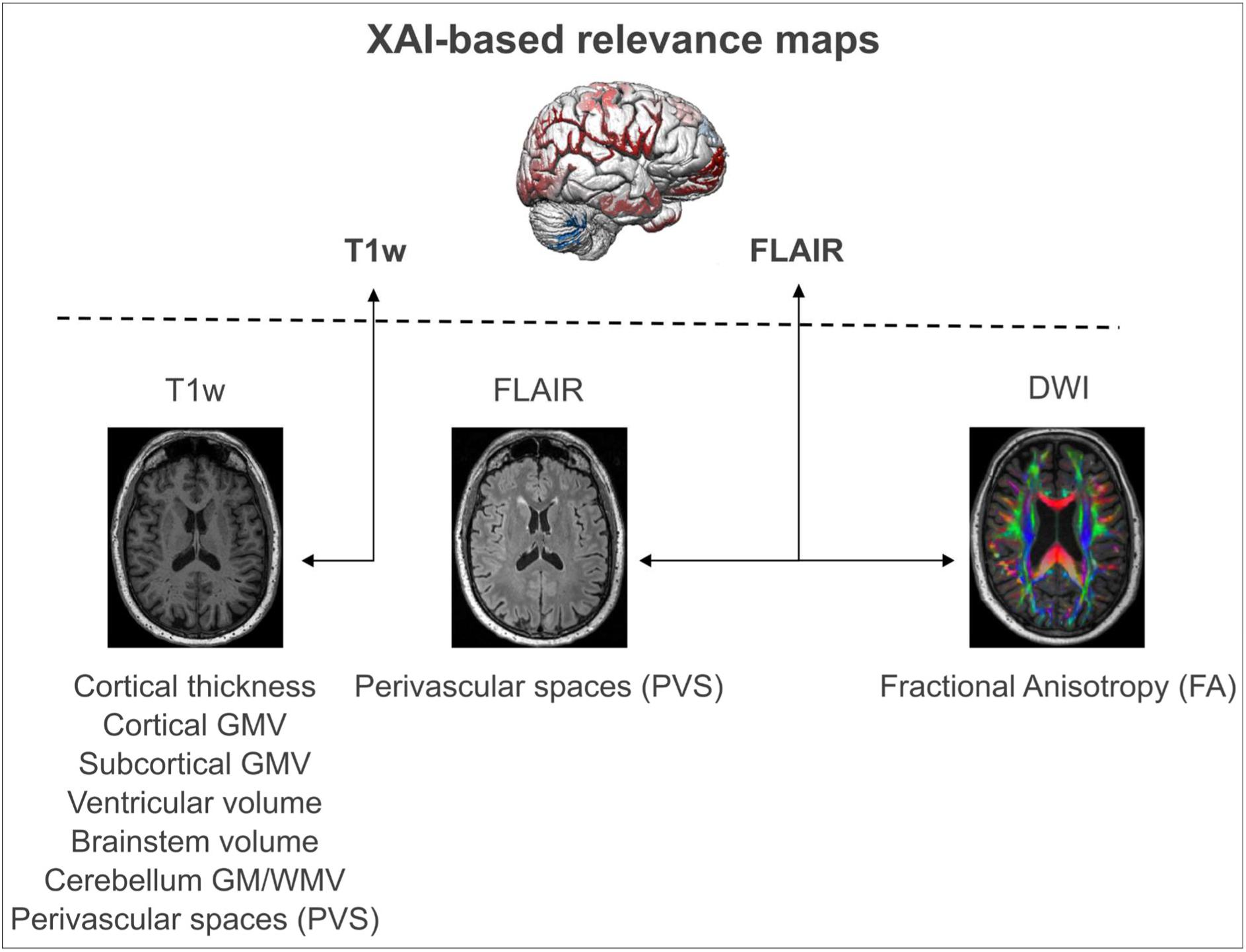
Relationship of aging-related imaging markers and deep learning-based relevance maps of brain age. Based on three imaging modalities (T1w, FLAIR, DWI) we computed eight different imaging markers. To test for their contribution to deep learning-based brain age predictions, we related these imaging markers to our computed LRP relevance scores. T1w-based imaging markers were related to T1w-based LRP relevance maps, FLAIR and DWI imaging markers were related to FLAIR-based LRP relevance maps. The contribution of white matter hyperintensities was already tested and reported in Hofmann et al. (2022). XAI: explainable artificial intelligence; GMV: gray matter volume

#### 2.5.1 Cortical thickness and volume

For 1855 participants, the regional mean cortical thickness (CT) and volume estimates were correlated with the sum of relevance scores, obtained from the T1 LRP relevance map, in 68 regions (see above) using Pearson’s correlation (coefficient *r)*. Bonferroni-correction was used to correct for multiple testing. The sum of relevance, rather than the mean, was used to account for differences in brain/region size (see <u>**Section 2.4.2**</u> above). By using the sum of relevance, we ensure that the scores are not influenced by variations in brain or brain region size.

#### 2.5.2 Subcortical, brainstem, cerebellar and ventricular volumes

For 1855 participants, we correlated subcortical, brainstem, cerebellar and ventricular volumes in each of their regions defined by various atlases (see above) with the corresponding sum of relevance scores, obtained from the T1 LRP heatmap using Pearson’s correlation. We used Bonferroni-correction to adjust for multiple significance testing.

#### 2.5.3 Perivascular spaces

For 1872 participants, we compared the average relevance of PVS voxels in the whole-brain and in areas in and around the basal ganglia (BG) with the average relevance within the brain using one-tailed paired t-tests. In contrast to the analyses above, here, we used the average relevance, since we were interested in whether the expected relevance was higher in PVS voxels than in the rest of the brain within the same participant. We conducted these tests for relevance values based on T1 and FLAIR sub-ensembles. For T1-relevance scores, we run the tests additionally with dilated PVS maps (dilation *d* in {1, 2} voxels), since PVS are fluid-filled and expected to have zero intensity in T1w imaging, which, in turn, would result in zero relevance scores (i.e., image values of zero do not contribute to a model’s prediction). We also tested a more conservative approach by comparing the relevance attributed to PVS to any other voxel which is predictive of higher-than-average brain age (i.e., with positive relevance), as used for WMH in Hofmann et al. (2022). To verify the relationship between PVS volumes and age, we additionally ran a Pearson correlation between the total number of PVS voxels in participants’ MRIs and their corresponding age at data acquisition.

#### 2.5.4 Fractional anisotropy

To test the relationship between FA values and LRP relevance scores, we performed region-based and voxel-wise analyses in 1872 participants. For the region-based analysis, we correlated the sum of the relevance scores from the FLAIR- and T1-based LRP heatmap with the average FA value for each JHU atlas region. Results were Bonferroni-corrected for multiple comparisons. For the voxel-wise analysis, we correlated the FA values with the relevance score of the FLAIR-based LRP heatmap for each voxel and across subjects in the MNI space.

## 3. Results

### 3.1 Cortical brain measures

Out of 68 Desikan-Killiany (DK; Desikan et al., 2006) brain regions (34 per hemisphere), 42 showed a statistically significant negative correlation between the relevance scores and CT, i.e., relevance for higher BA was associated with lower CT (mean *r* = -.20, *r*-range = [-.44, - .09]). The strongest negative correlation was present in the inferior-parietal area in the right hemisphere (*r* = -.44, p < 8.1e-89). Only one area, the right pars orbitalis, showed a positive relationship, i.e., relevance for higher brain age was associated with higher CT (*r* = .11, p < .00024; **Fig.2**).

**Fig. 2.**
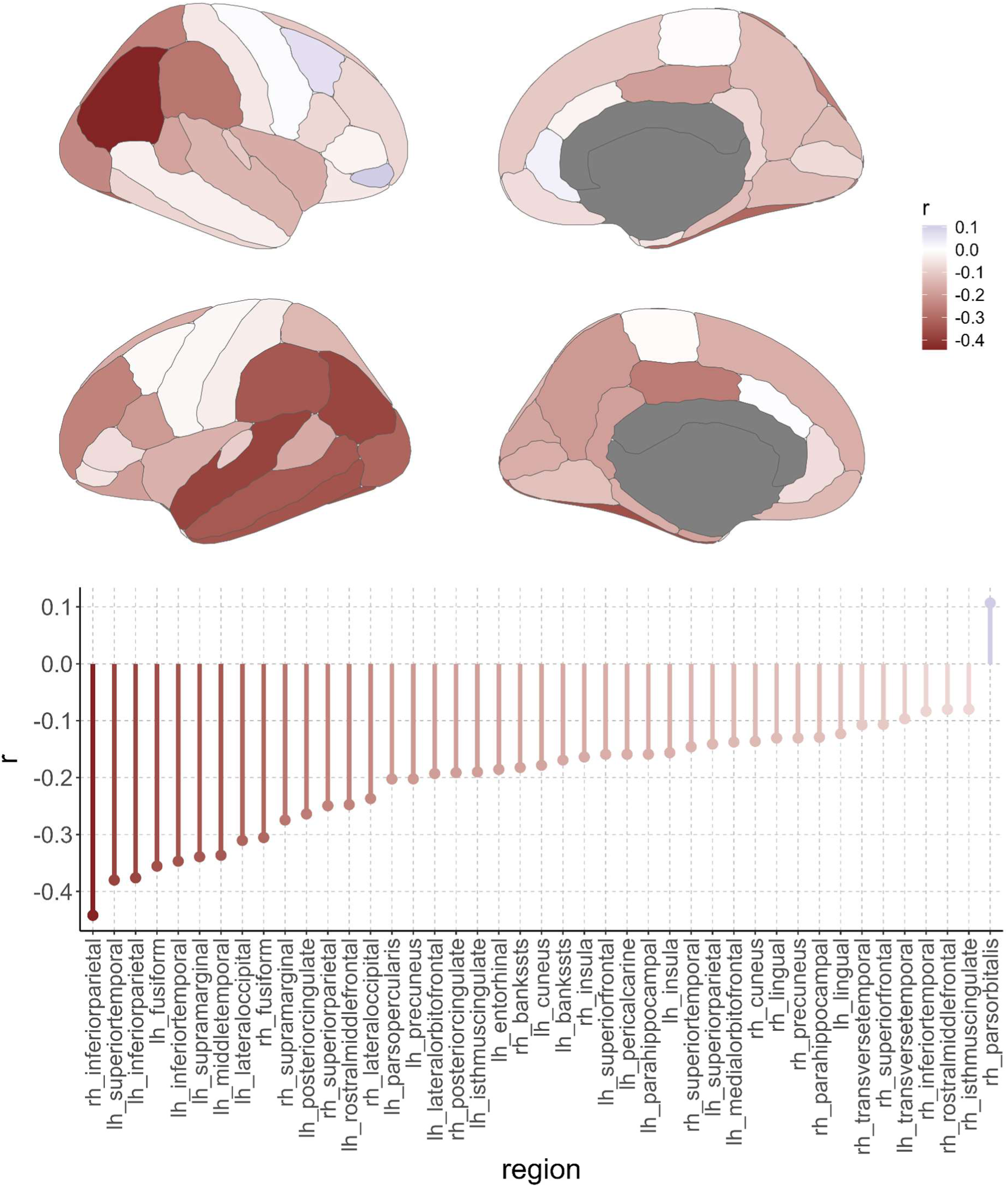
Relationship of cortical thickness with relevance scores from deep learning-based brain age predictions across the full lifespan in the LIFE-Adult study (n =1855). *Top panel*: Voxel-wise correlation results for the association between cortical thickness and relevance scores, color represents Pearson’s correlation coefficient. *Bottom panel*: Region-wise results for Bonferroni-corrected significant associations between cortical thickness and relevance scores (43 out of 68 regions defined by the DK atlas).

Out of 68 DK brain regions, 44 showed a statistically significant negative relationship between the LRP-based relevance scores and GMV (Bonferroni corrected, mean *r* = -. 22, *r*-range = [-.40, -.08]). The strongest negative correlation was present in the inferior temporal gyrus in the left hemisphere (*r* = -.40, p < 1.5e-70). Four areas showed a statistically significant positive relationship (Bonferroni corrected, mean *r* = .11, *r*-range = [.10, .11]). The strongest positive correlation was present in the middle temporal gyrus in the right hemisphere (*r* = .11, *p* < .0001; **Fig.2**).

**Fig. 2:**
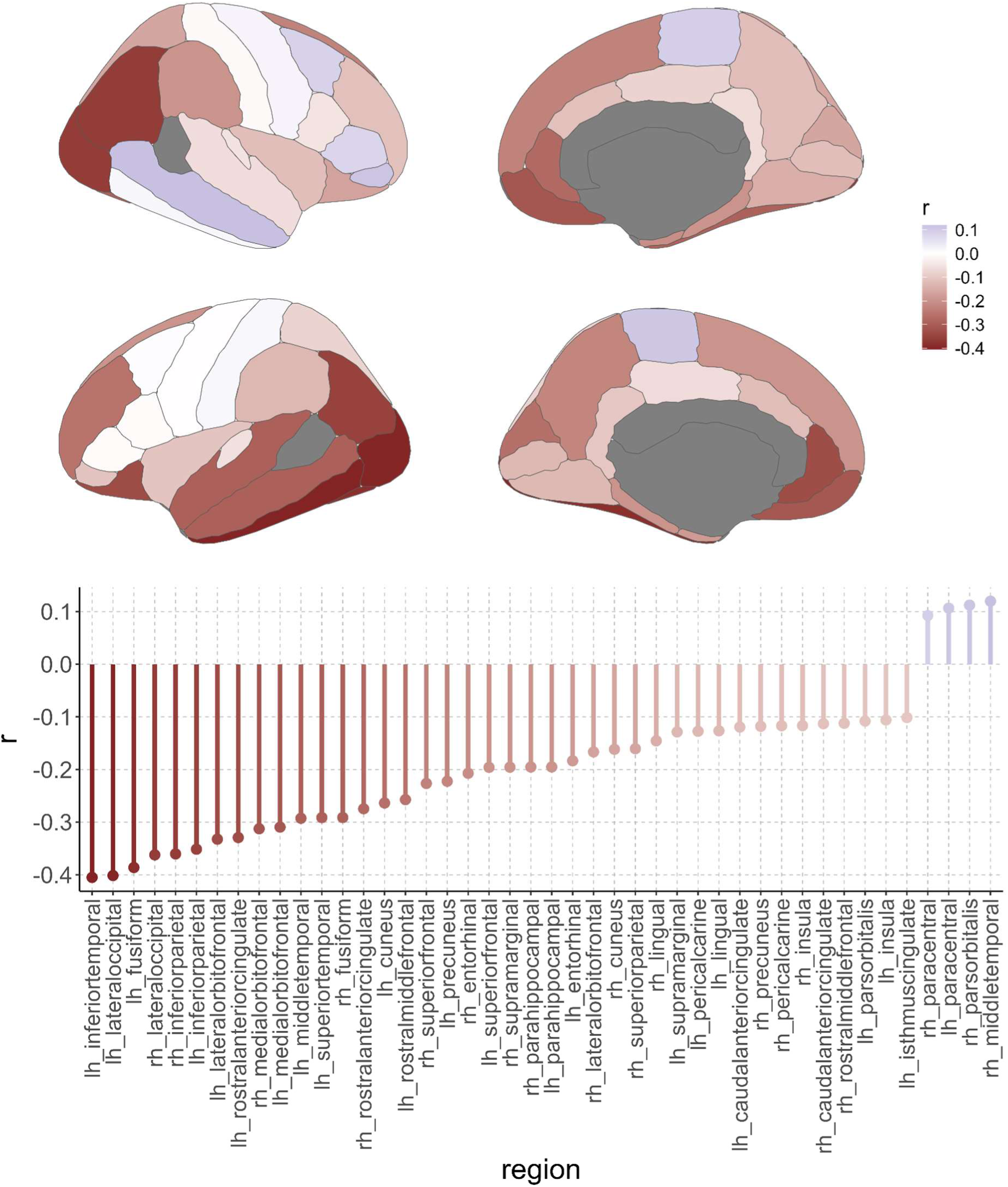
Relationship of cortical gray matter volume with relevance scores from deep learning-based brain age predictions across the full lifespan in the LIFE-Adult study (n = 1855). *Top panel*: Voxel-wise correlation results for the association between cortical gray volume and relevance scores, color represents Pearson’s correlation coefficient. *Bottom panel*: Region-wise results for Bonferroni-corrected significant associations between cortical gray volume and relevance scores (48 out of 68 regions defined by the DK atlas).

### 3.2 Subcortical brain measures

Out of 25 subcortical regions, including brainstem and cerebellum from the FreeSurfer *aseg* atlas, 19 (all regions except of the ventricles and cerebrospinal fluid; CSF) showed a statistically significant negative relationship between the LRP-based relevance scores and GMV (Bonferroni corrected, mean *r* = -.28, *r*-range = [-.52, -.14]). The strongest negative correlation was present with the left cerebellar GMV (*r* = -.52, p < 8.9e-141). Five regions (all of them ventricles or CSF) showed a statistically significant positive relationship (Bonferroni corrected, mean *r* = .44, *r*-range = [.26, .69]). The strongest positive correlation was present in the right lateral ventricle (*r* = .69, p < .2.5e-284; **Fig.3**).

**Fig. 3:**
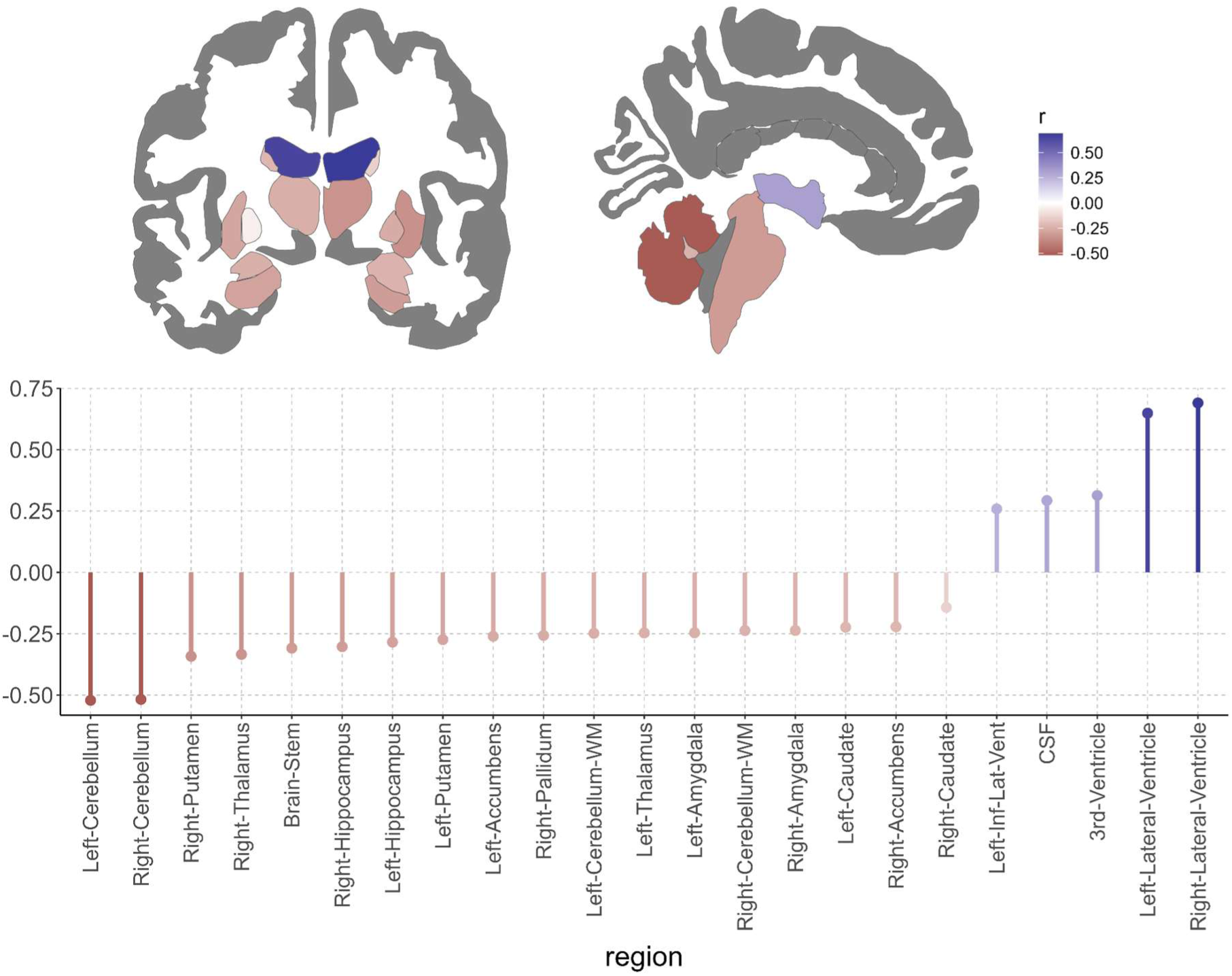
Relationship of subcortical, cerebellar gray matter, brainstem and ventricle volume with relevance scores from deep learning-based brain age predictions across the full lifespan in the LIFE-Adult study (n = 1855). *Top panel*: Voxel-wise correlation results for the association of subcortical, cerebellar gray matter, brainstem and ventricle volume with relevance scores, color represents Pearson’s correlation coefficient. *Bottom panel*: Region-wise results for Bonferroni-corrected significant associations of subcortical, cerebellar gray matter, brainstem and ventricle volume with relevance scores (23 out of 25 regions defined by the FreeSurfer *aseg* atlas).

### 3.3 Cerebral small vessel disease (cSVD) and white matter-related measures

#### 3.3.1 Perivascular spaces (PVS)

In line with previous literature, we found that the number and magnitude of PVS increases with age both in the whole brain (*r* = .403, p < 5.3e-74) and areas in and around the basal ganglia (*r* = .16, p < 2.9e-12; see **Fig. S1** in the Supplementary materials).

There was no significant difference between relevance scores extracted from the T1 sub-ensemble for both PVS maps with no dilation (*d* = 0) and with dilations (*d* in {1, 2}) and the average relevance scores across the brain (one-tailed paired t-tests, no dilation: t*_d_*_=0_(1871) = -1.17, p*_d_*_=0_ = .88, **Fig.4a,** also see **Fig. S4a** in Supplementary materials; with dilation, t-test: t*_d_*_=1_(1871) = -0.93, p*_d_*_=1_ = .82, **Fig. S4b**; t*_d_*_=2_(1871) = -0.058, p*_d_*_=2_ = 0.52, **Fig. S4c**). This was also the case when only considering PVS around BG (one-tailed paired t-test*_d_*_=1_, t(1871) = -3.85, p = 0.99; t-test*_d_*_=2_, t(1871) = -5.98, p = 1).

Relevance scores extracted from the FLAIR sub-ensemble were significantly higher in PVS than the relevance score averaged over the whole brain (one-tailed paired t-test, t(1871) = 14.82, p < 2.2e-47, one-tailed; **Fig.4b**), however, when using the conservative approach comparing average PVS relevance to the average positive relevance in the whole brain only, no significant difference was found (t(1871) = -121.14, p = 1, one-tailed, **Fig. S2b** in Supplementary materials). Also, when only considering PVS around BG there was no significant difference (paired t-test, t(1871) = 0.284, p = 0.388, one-tailed). In Figure 4c, we show the difference between relevance values in WMH compared to overall positive relevance for comparison, taken from Hofmann et al. (2022).

**Fig. 4:**
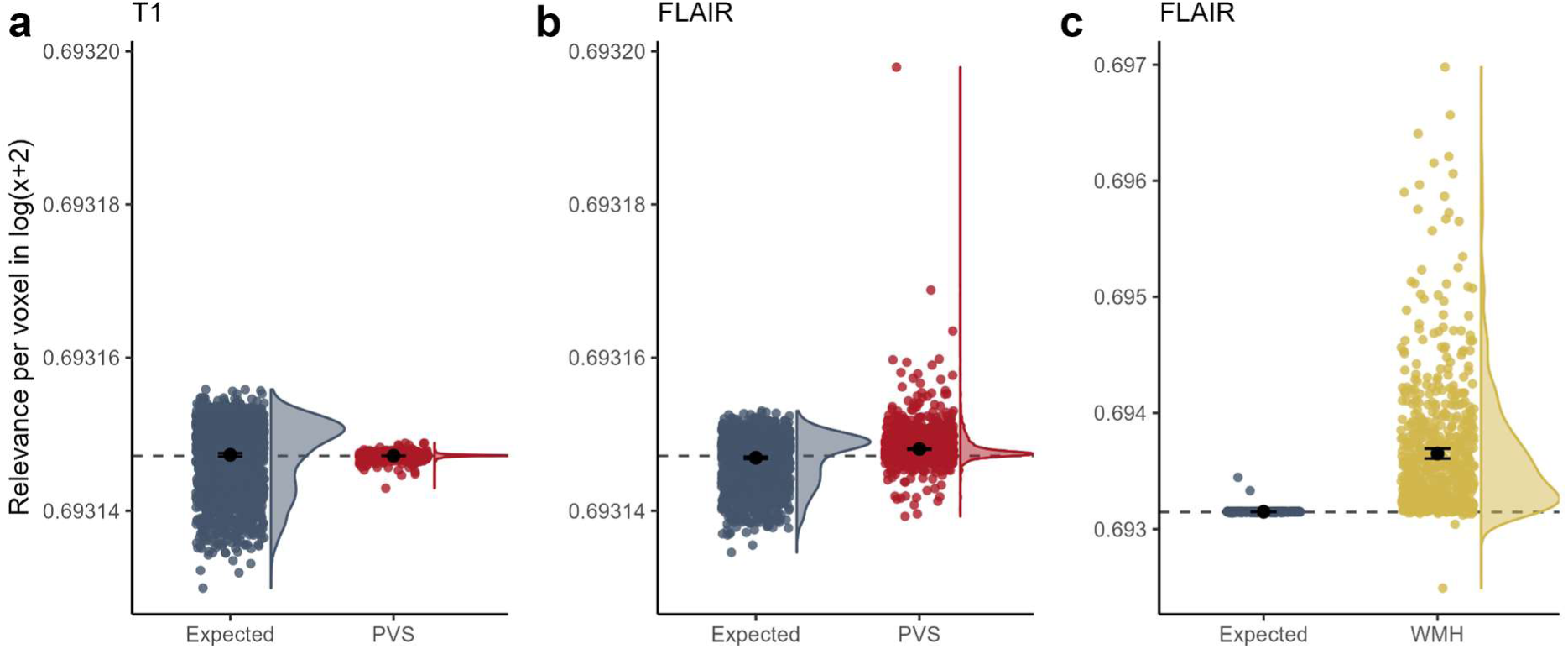
Average relevance in voxels classified as perivascular spaces (PVS). (**a**) Average relevance in PVS voxels (in red) and expected whole-brain relevance (in blue) in LRP maps from the T1 sub-ensemble (n = 1872). (**b**) Average relevance in PVS voxels (in red; not dilated) and expected relevance (in blue) in LRP maps from the FLAIR sub-ensemble. (**c**) Average relevance in white matter hyperintensity (WMH) voxels (in yellow) and expected whole-brain relevance (in blue) in LRP maps from the FLAIR sub-ensemble (reproduced from Hofmann et al., 2022). Relevance scores are shown in log(x+2) scale for better visual representation. Data points were horizontally spread (using *geom_jitter* in *ggplot2)* to enhance visibility.

Upon visual inspection of some individuals with high numbers of PVS, we did not see a systematic correspondence of relevance maps and PVS segmentations, while only some individual PVS were overlapping with higher relevance scores (see **Fig. S3** in Supplementary materials).

#### 3.3.2 Fractional anisotropy

We performed a region-based and a voxel-wise correlation analysis, investigating the relationship between LRP-based relevance scores and diffusion tensor imaging (DTI)-based fractional anisotropy (FA). For the region-based analysis that related relevance scores from the FLAIR sub-ensemble to FA, we found that in 23 out of 48 regions of the DTI-based WM atlas JHU higher FA values were associated with lower relevance values (Bonferroni corrected, mean *r* = -.13, *r*-range = [-.25, -.07]). The highest negative correlation was present in the left superior fronto-occipital fasciculus (*r* = -.25, p < 1.5e-33). For one region, the left superior cerebellar peduncle, lower FA values were associated with lower relevance values (*r* = .07, p < .01; **Fig.5**). The voxel-wise correlation analysis per subject revealed a predominantly negative correlation between FA values and relevance scores (similar to the region-based analysis), with higher FA values associated with lower relevance scores, i.e., lower BA (mean *r* = -.02, *r*-range = [-.19, .09]).

**Fig. 5:**
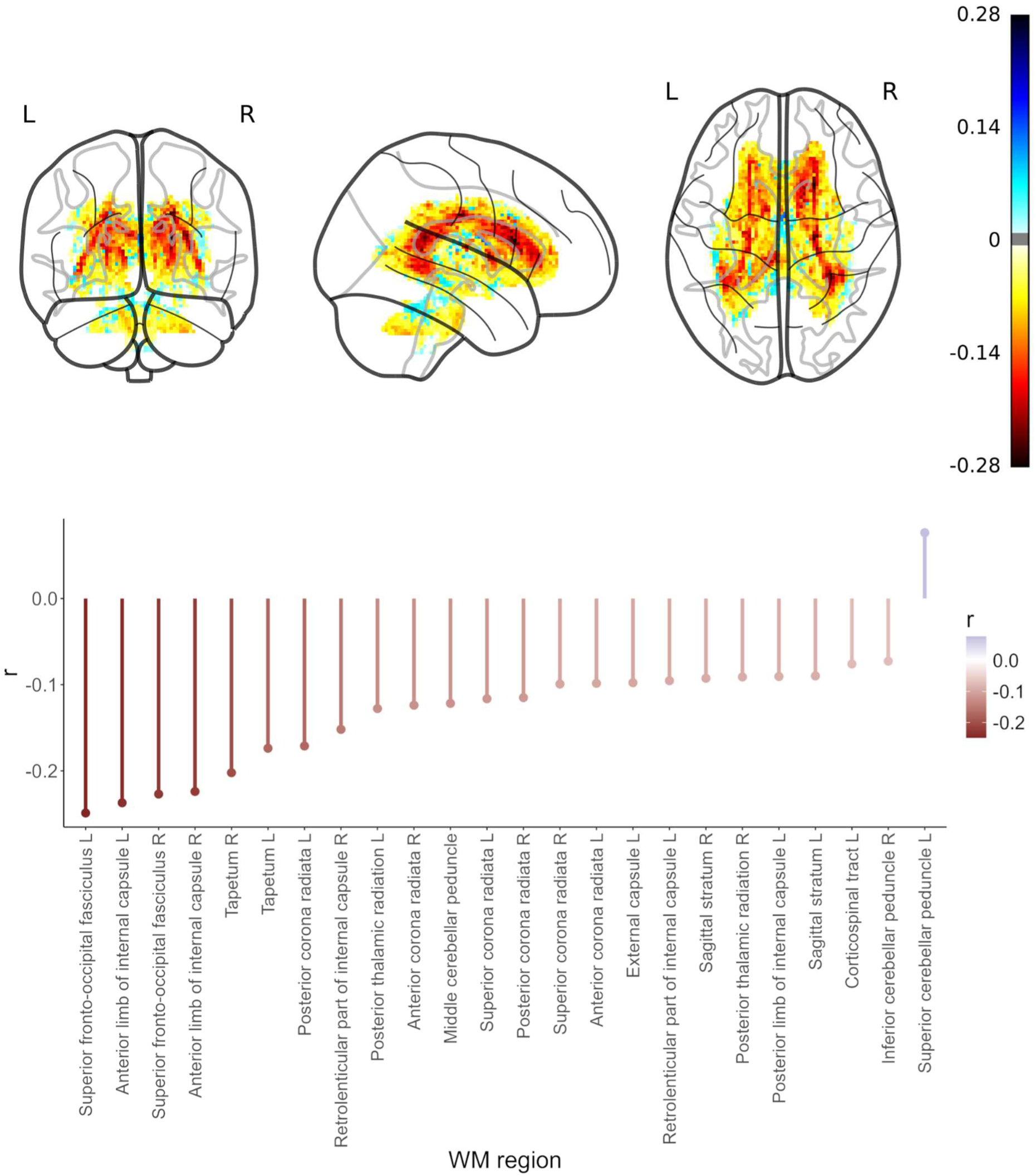
Relationship of fractional anisotropy (FA) with relevance scores from deep learning-based brain age predictions across the full lifespan in the LIFE-Adult study (n = 1855). *Top panel*: Voxel-wise correlation results for the association between FA and relevance scores, color represents Pearson’s correlation coefficient. *Bottom panel*: Region-wise results for Bonferroni-corrected significant associations between FA and relevance scores from the FLAIR sub-ensemble (24 out of 48 regions defined by the JHU atlas). Region-wise results for associations between FA and relevance scores from the T1 sub-ensemble are shown in **Fig. S5** in **Supplementary materials**.

For the region-based analysis that related relevance scores from the T1 sub-ensemble to FA, we found that in 29 out of 48 JHU regions higher FA values were associated with lower relevance values (Bonferroni corrected, mean *r* = -.15, *r*-range = [-.31, -.05]), similar to the relevance scores from FLAIR sub-ensemble. The highest negative correlation was present in the right tapetum (*r* = -.31, p < 2.5e-102). In four regions, lower FA values were associated with lower relevance values (mean *r* = .06, *r*-range = [.05, .09]; **Fig. S5**).

## Discussion

In this work, we showed that DNN-based BA predictions were correlated with widespread differences in common neuroimaging features of aging. In particular, higher XAI-based relevance was associated with lower cortical thickness and volume in frontal-temporal and parietal brain regions. The highest correlation of all features was found between ventricular volume and ventricular relevance. Higher relevance in limbic and basal ganglia regions was also associated with lower GMV. Interestingly, of all non-cerebral features, higher XAI-based relevance in the cerebellum was most strongly associated with lower cerebellar GMV. Regarding WM differences, we found that higher relevance in frontal-occipital fasciculus and anterior limb of the internal capsule were strongly negatively related to directedness of WM microstructure in these regions. Contrary to our expectations, PVS were not reliably associated with higher relevance, that is, the DNN models trained on T1 or FLAIR did not consistently consider PVS relevant for predicting BA. Taken together, our method highlighted known imaging features of brain aging, but also highlighted some less considered regions such as the cerebellum.

### 4.1 Relevance of cortical and non-cortical structures for brain age

Our finding of a frontal-temporal parietal pattern of negative associations between relevance and cortical GM features is in line with studies showing linear age associations in superior and inferior frontal gyri, parietal cortex and superior parts of the temporal lobe (Aycheh et al., 2018; Cox et al., 2021; Fjell et al., 2009; Fjell & Walhovd, 2010). Against our expectation, we found some cortical features in the frontal and temporal lobe to be positively associated with relevance scores. Overall, these associations were weaker than the negative associations and might reflect spurious associations due to missegmentations of the pial surface in FreeSurfer. Regarding non-cortical structures, our results of strongest association between ventricular volume and XAI-based relevance confirmed previous findings of the age-related expansion of the ventricular system. We found the largest association for lateral ventricles which showed the 2nd strongest effect of 4.4 % longitudinal annual average change in (Fjell et al., 2013). We also found that cerebellar GMV among all non-cerebral GM features was most strongly associated with relevance, which was surprising given that cerebellum GMV showed a lower association with age than, for instance, nucleus accumbens and hippocampus in previous literature (Fjell et al., 2013). FreeSurfer, which was designed to segment the cerebral cortex, might perform suboptimally when determining the GMV of the cerebellum, while the DL model takes an unbiased approach in every region uniquely relying on signal intensity. However, as a recent review suggests, the cerebellum might have a widely underappreciated role in brain aging both structurally and cognitively (Bernard, 2022). In future research it would be worth exploring the cognitive consequences in participants with particularly high BA relevance scores in cerebellar tissue.

### 4.2 Relevance of perivascular spaces for brain age

We did not find PVS to be consistently considered by the DNNs for brain age prediction. Compared to WMH, PVS, another common imaging marker of cerebral small vessel disease (cSVD), only showed a weak, yet significant increase of FLAIR-based relevance in whole-brain PVS (but not basal-ganglia PVS), and no difference for T1-based relevance maps. Visually we did not see a systematic overlap of relevance values with the PVS segmentation when inspecting individuals with large visible PVS. We therefore speculate that the increased FLAIR-based relevance might have been confounded by two factors: First, some of the deep WM PVS considered were close to the cortical surface, which showed consistently higher relevance values both based on T1 and FLAIR sub-ensembles (see Hofmann et al., 2022). Second, WMH are known to form around PVS (Wardlaw et al., 2020), which could result in higher FLAIR relevance attributed to PVS. From a mathematical point of view, PVS in T1w-images should be surrounded by higher relevance values rather than overlapping with them, given that the BA model considers them as relevant. This is because only non-zero image intensity values can contribute to the model estimates of BA and PVS or other fluid containing spaces usually have zero-intensity values in T1w images. We therefore dilated the PVS masks to detect surrounding relevance scores in T1-based relevance maps. However, in contrast to FLAIR-based maps for T1-based relevance maps, we did not find any difference between PVS and all other voxels, even though PVS are commonly detected on T1w images (Duering et al., 2023). This was the case no matter whether original or dilated PVS maps were used, indicating that the shape or spatial features of this structure were not taken into account by the DNNs. For areas in and around the basal ganglia, we found that the association between age and PVS volume was smaller as compared to whole-brain PVS (**Fig. S1**), therefore, we can also expect a smaller relationship between BA relevance scores and PVS in these regions.

As the two cSVD imaging markers WMH and PVS tend to co-occur (Wardlaw et al., 2020), the lack of consideration of the DNNs for PVS might indicate that WMH and PVS are not sufficiently distinct (i.e., their appearance might be topologically collinear) to provide additional information to the model. Previous research also demonstrated that the relationship between PVS and age is modulated by other features such as intracranial volume and hypertension (Huang et al., 2021). Although we did find a robust correlation between the total number of PVS voxels and age (**Fig. S1**), the relationship with PVS may be too subtle (Barisano et al., 2021; Lynch et al., 2023) to be captured by our BA model, when compared to more salient aging processes in the brain.

### 4.3 Relevance of white matter structures for brain age

We found that higher relevance in periventricular frontal and occipital WM was most strongly negatively associated with attributed XAI-based relevance. This finding reflects the strong association of aging with WMH, and their highest prevalence in periventricular WM (De Leeuw et al., 2001). We showed previously that the average relevance in WMH was significantly larger than the average positive relevance outside WMH (Hofmann et al., 2022). Here, we used relevance based on DNN-models trained on FLAIR, which reflect signal intensity differences related to WMH. Diffusion-related imaging metrics are known to be severely altered in WMH, displaying reduced FA and increased mean diffusivity which reflects increased water content, and in severe cases damaged fiber tracts in these regions (Wardlaw et al., 2015). With our DNN models trained on FLAIR, which is primarily susceptible to tissue water content, we could not quantify the contribution of other subthreshold WM differences which might appear from a model trained on diffusion-weighted imaging. Interestingly, we also found a relationship between FA values and relevance scores obtained from the T1 sub-ensemble in approximately 69% of the WM regions. The regions with the highest negative correlation with FA showed a strong overlap between FLAIR and T1-based relevance maps. While it is known that T1 values are altered in severe WHMs (Maniega et al., 2015), our results suggest that the T1 sub-ensemble might be capable of detecting more subtle changes in T1 intensities to predict brain aging.

### 4.4 Limitations

So far, we could not provide an indication to which extent further unknown imaging features contribute to the BA estimation for the 3D-DNNs. We did not directly compare FA values with relevance based on the same imaging modality, and the association with FLAIR-based relevance might have underestimated the association with age as FLAIR is not susceptible to diffusion-based differences except the impact of tissue water. In future work, an additional sub-ensemble could be attached to the BA model architecture where DTI is utilized for model training and, subsequently, XAI-based relevance maps can be computed directly from the DTI model branch.

Also, relevance maps of LRP should be carefully evaluated. First, they could be selectively interpreted to confirm *a priori* assumptions leading to a confirmation bias (Ghassemi et al., 2021). We therefore chose to report results across the whole brain, avoiding selective ROIs. Second, not all relevance values within a map must be meaningful with respect to the predicted (target) variable, hence, one must assume a certain level of noise in these maps. This can be illustrated by untrained neural networks that are often sensitive towards strong image contrasts resulting in spurious informative saliency maps (Adebayo et al., 2020). We therefore chose group-level statistics to average out noise effects over participants. Particularly for within-subject analyses, one could consider recently promoted techniques for DNN-based modeling that mitigate potential confounding effects resulting from preprocessing steps such as skull stripping and image registration (Tinauer et al., 2022). Third, LRP (and similar XAI approaches) highlight only voxel-wise contributions to the model estimates, while underlying relationships between voxels can only be inferred with domain knowledge. There are new variants of LRP that allow to extract higher order structure (Eberle et al., 2022; Schnake et al., 2022) or underlying relevant concepts (Achtibat et al., 2023; here, e.g., this could be interacting structural changes of two brain regions); hence, it would be a promising path of research to employ Concept Relevance Propagation (see Tinauer et al., 2024).

## Conclusion

Taken together, we show that highly accurate DNN-based multi-modal BA predictions are partially driven by morphometric changes in the cortex, subcortex and WM. While most age-related cortical thinning, subcortical GM atrophy, ventricular expansion, and WMH appearance have been investigated earlier, we showed that these relationships can be discovered by end-to-end models, that is, these models are not explicitly trained on these features but trained on relatively unprocessed MR images to estimate age. At the same time, also less considered regions such as cerebellum are highlighted by this data-driven approach. End-to-end DNNs generally have higher predictive accuracy compared to linear models or other classic machine learning approaches that are trained on pre-selected features (e.g., see the brain-age competition PAC 2019). Here, we have provided evidence that DNNs consider these biologically relevant brain features. However, their high prediction performance was only partially explained by these features. In future work, our explainable A.I. pipeline could help to identify unknown brain changes, which could be further enhanced by high-resolution imaging.

## Data and Code Availability

Due to potential identifiability of individuals from demographic and medical information, we cannot share the processed data used in this study. Raw data of the LIFE-Adult cohort can be requested via the LIFE study center (https://ldp.life.uni-leipzig.de/).

All code is in https://github.com/SHEscher/RelevanceRelated.

## Author Contributions

SMH: Conceptualization, Methodology, Software, Validation, Formal analysis, Investigation, Data curation, Writing—original draft, Writing—review & editing, Visualization, Project administration

OG: Conceptualization, Methodology, Software, Validation, Formal analysis, Investigation, Data curation, Writing—original draft, Writing—review & editing, Visualization

NS: Writing—review & editing, Supervision, Funding acquisition

KRM: Writing—review & editing, Supervision

ML: Resources, Funding acquisition

AV: Resources, Funding acquisition

MG: Writing—review & editing, Supervision

AVW: Writing—review & editing, Supervision

FB: Conceptualization, Methodology, Software, Validation, Formal analysis, Writing—original draft, Writing—review & editing, Visualization, Funding acquisition

## Acknowledgements

We thank the participants of the LIFE-Adult cohort for their participation.

## Supplementary materials

**Fig. S1:**
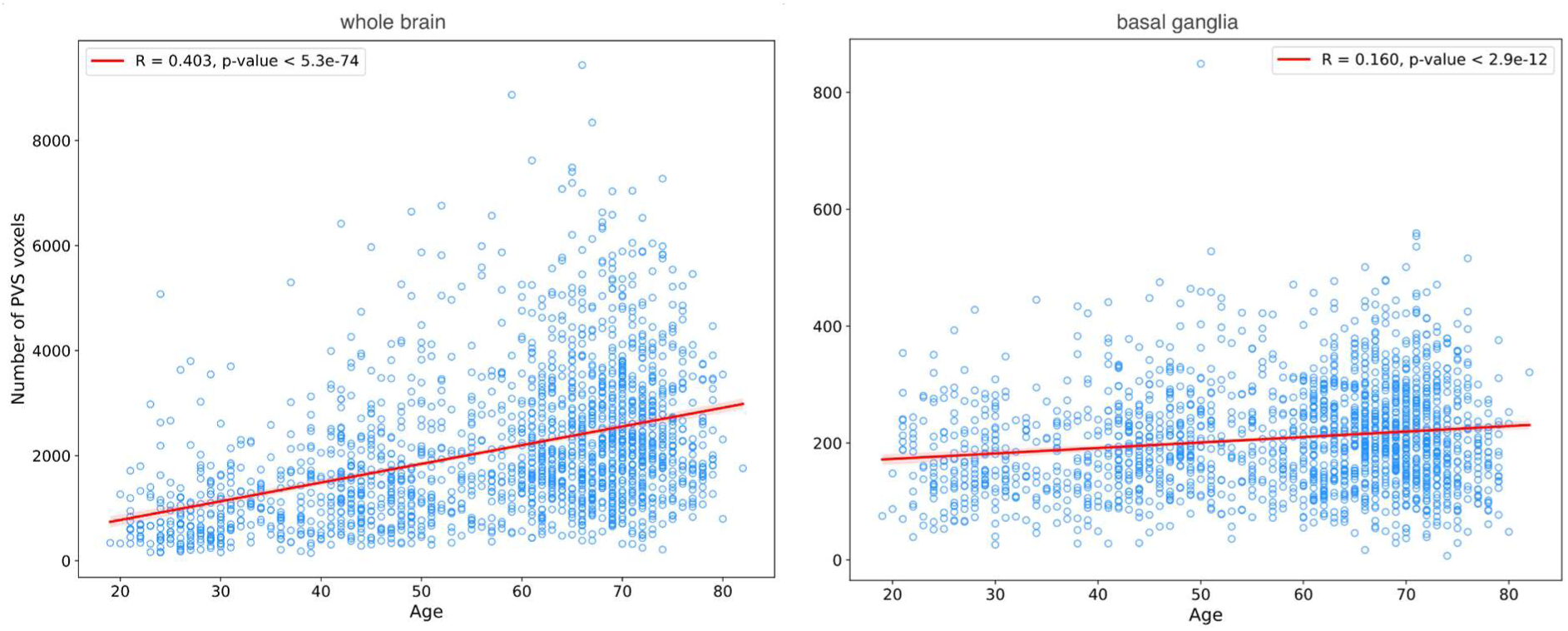
Relationship between age and perivascular spaces (PVS) in the whole brain (*left*) and in the area in and around the basal ganglia (*right panel*). PVS segmentation was done using SHIVA-PVS toolbox (Boutinaud et al., 2021; https://github.com/pboutinaud/SHIVA_PVS; accessed in March 2024). The basal ganglia was defined by the ATAG atlas (https://www.nitrc.org/projects/atag), and dilated by 5 voxels to include deep white matter areas surrounding the basal ganglia. *Red line*: Pearson correlation *R* between age and number of PVS voxels across participants.

**Fig. S2:**
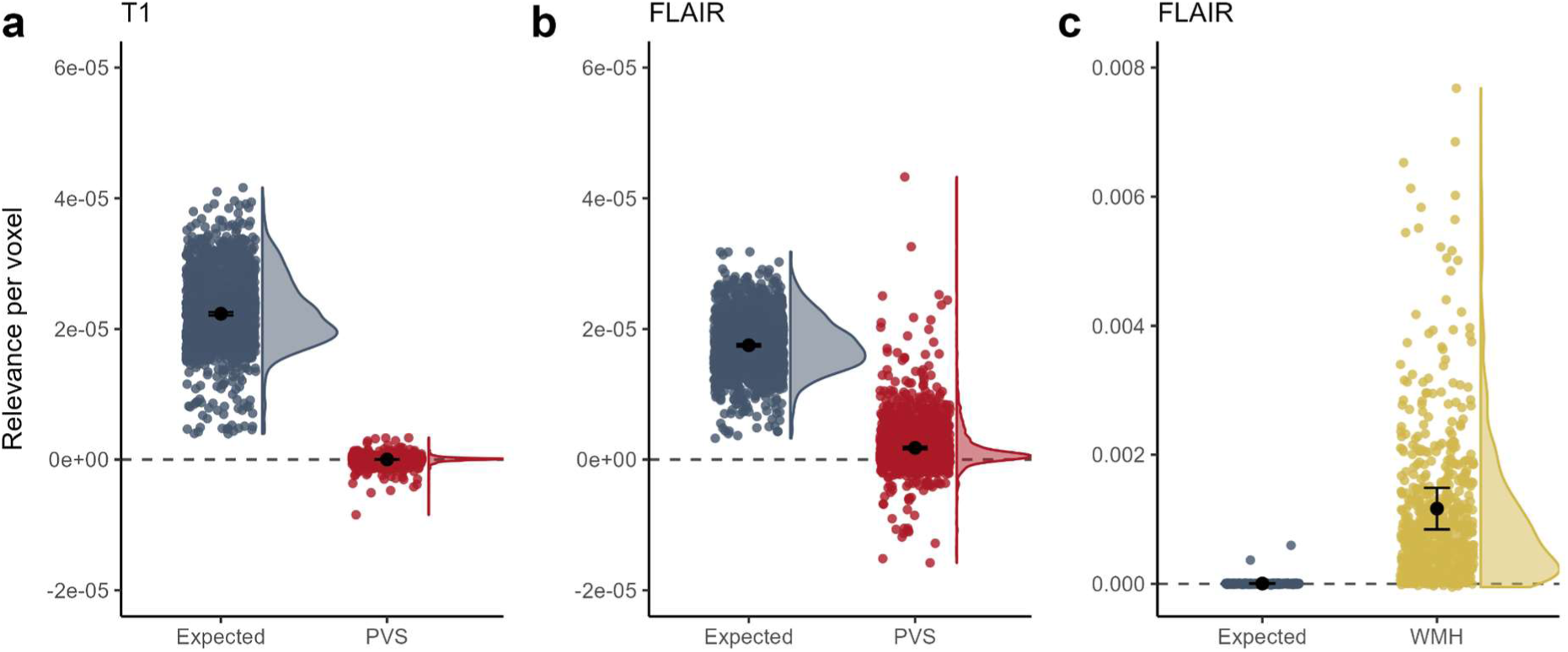
Average relevance in voxels of perivascular spaces (PVS) compared to average relevance of voxels with positive relevance. (**a**) Average relevance in PVS voxels (in red) and expected relevance (in blue) for LRP-heatmap of the T1 sub-ensemble (n = 1855). (**b**) Average relevance in PVS voxels (in red) and expected relevance (in blue) for LRP-heatmap of the FLAIR sub-ensemble. (**c**) Average relevance in WMH voxels (in yellow) and expected relevance (in blue) for LRP-heatmap of the FLAIR sub-ensemble (reproduced from Hofmann et al. (2022). Here, expected relevance is defined as the average relevance score of all positive relevance voxels in the whole brain for the corresponding participant. This is a more conservative comparison level, as compared to all relevance voxels as a baseline. Data points were plotted using *geom_jitter* in *ggplot2* to enhance visibility.

**Fig. S3:**
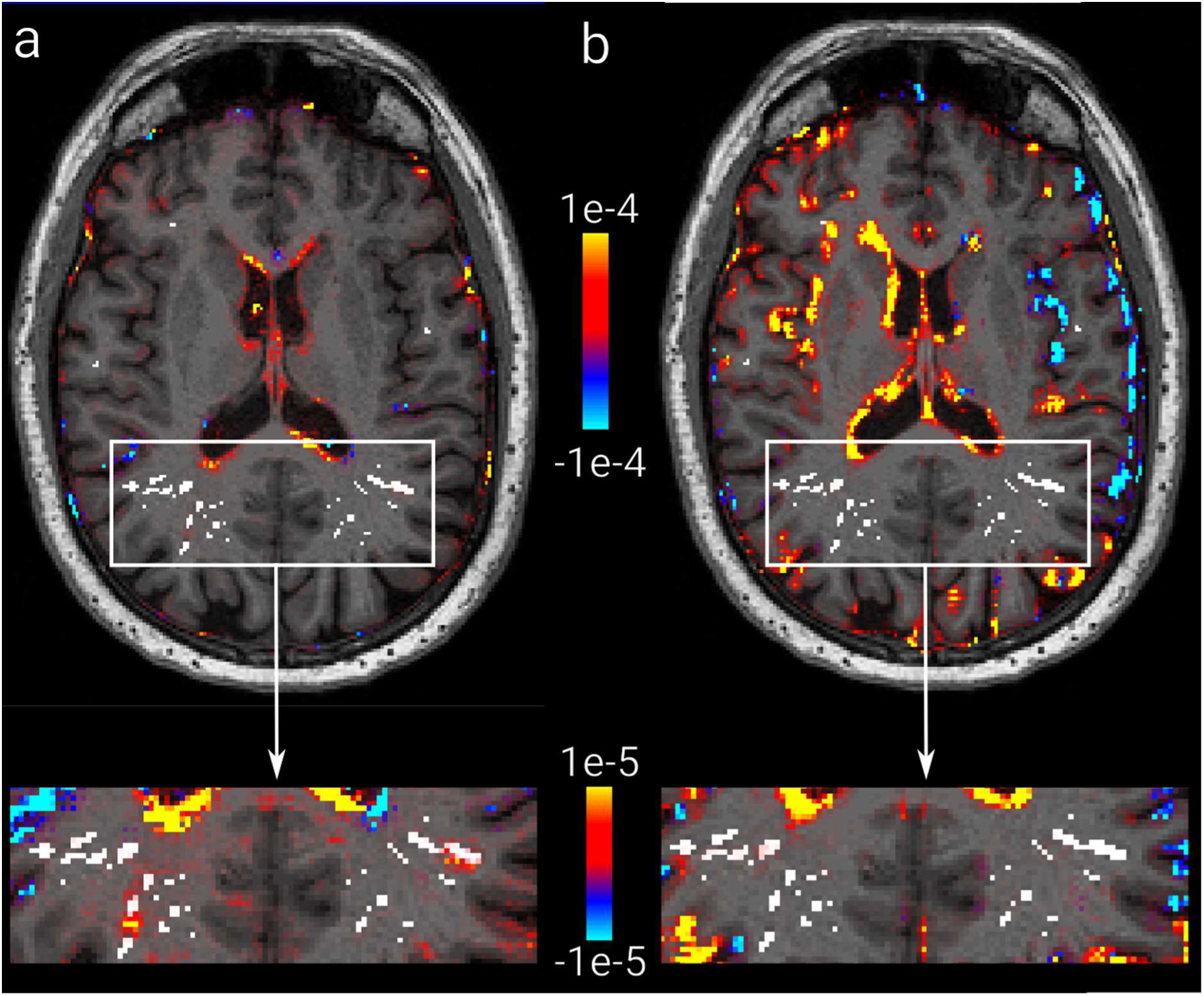
Perivascular space (PVS) segmentation and brain age relevance in a 65-year old male. **A**: PVS segmentation (white) and relevance values from LRP based on the T1 sub-ensemble overlaid on the corresponding T1w image. **B**: PVS segmentation (white), relevance values from LRP based on FLAIR sub-ensemble. (upper panels: red/blue: positive/negative relevance thresholded at 10^-4^, lower panels: relevance thresholded at 10^-5^)

**Fig. S4:**
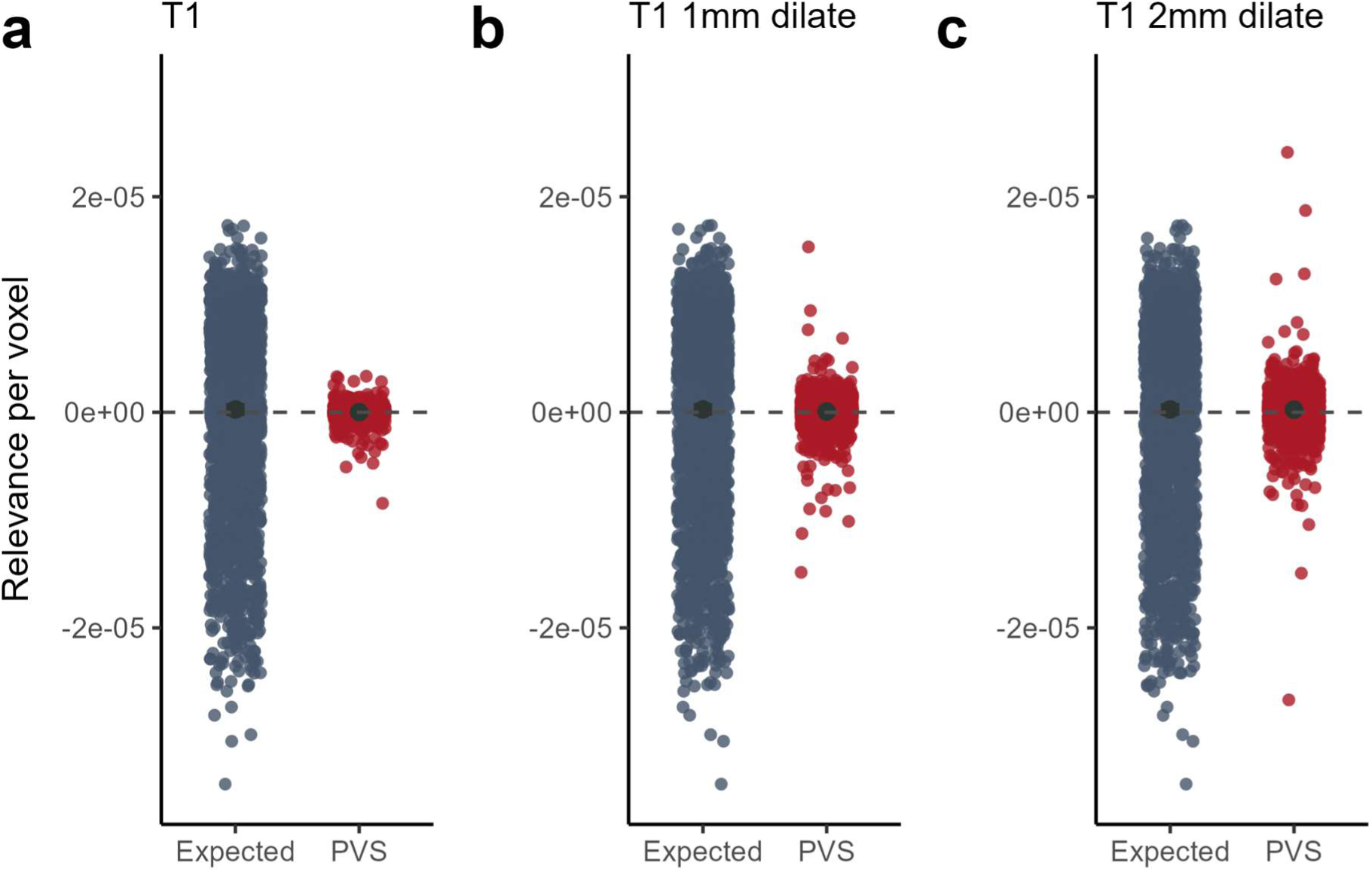
Effect of dilation on T1 sub-ensemble based relevance values of PVS. (**a**) Average relevance in PVS voxels (in red) and expected relevance (in blue) for LRP-heatmap of the T1 sub-ensemble (n = 1855). (**b**) Average relevance in PVS voxels (in red) and expected relevance (in blue) for LRP-heatmap of the T1 sub-ensemble with 1 mm dilation. (**c**) Average relevance in PVS voxels (in red) and expected relevance (in blue) for LRP-heatmap of the T1 sub-ensemble with 2 mm dilation. For none of the three, the relevance in PVS is statistically significantly greater than the expected relevance. Data points were plotted adding small random variation to the location of each datapoint to enhance visibility (using *geom_jitter* in *ggplot2*).

**Fig. S5:**
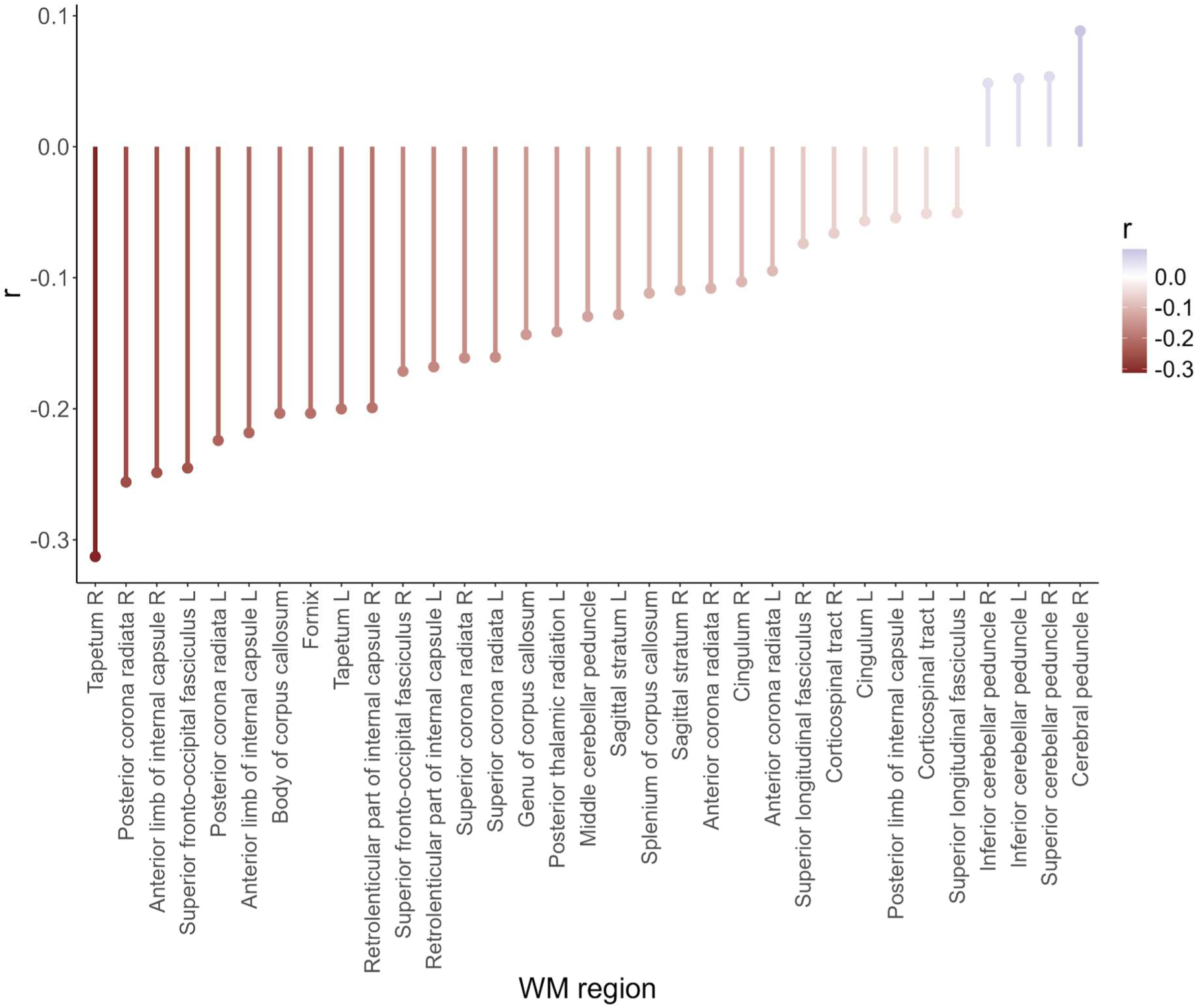
Relationship of regional fractional anisotropy (FA) with relevance scores obtained from T1 sub-ensemble LRP heatmap across the full lifespan in the LIFE-Adult study (n = 1855). Only Bonferroni-corrected statistically significant associations are shown (33 out of 48 regions defined by the JHU atlas).

## References

Achtibat, R., Dreyer, M., Eisenbraun, I., Bosse, S., Wiegand, T., Samek, W., & Lapuschkin, S. (2023). From attribution maps to human-understandable explanations through Concept Relevance Propagation. Nature Machine Intelligence, 5(9), 1006–1019. 10.1038/s42256-023-00711-8

Adebayo, J., Gilmer, J., Muelly, M., Goodfellow, I., Hardt, M., & Kim, B. (2020). Sanity Checks for Saliency Maps. http://arxiv.org/abs/1810.03292

Avants, B. B., Epstein, C. L., Grossman, M., & Gee, J. C. (2008). Symmetric diffeomorphic image registration with cross-correlation: Evaluating automated labeling of elderly and neurodegenerative brain. Medical Image Analysis, 12(1), 26–41. 10.1016/j.media.2007.06.004

Aycheh, H. M., Seong, J.-K., Shin, J.-H., Na, D. L., Kang, B., Seo, S. W., & Sohn, K.-A. (2018). Biological Brain Age Prediction Using Cortical Thickness Data: A Large Scale Cohort Study. Frontiers in Aging Neuroscience, 10, 252. 10.3389/fnagi.2018.00252

Bach, S., Binder, A., Montavon, G., Klauschen, F., Müller, K.-R., & Samek, W. (2015). On Pixel-Wise Explanations for Non-Linear Classifier Decisions by Layer-Wise Relevance Propagation. PLOS ONE, 10(7), e0130140. 10.1371/journal.pone.0130140

Bacon, F. (1620). Novum Organum.

Ball, G., Kelly, C. E., Beare, R., & Seal, M. L. (2021). Individual variation underlying brain age estimates in typical development. NeuroImage, 235, 118036. 10.1016/j.neuroimage.2021.118036

Barisano, G., Sheikh-Bahaei, N., Law, M., Toga, A. W., & Sepehrband, F. (2021). Body mass index, time of day and genetics affect perivascular spaces in the white matter. Journal of Cerebral Blood Flow & Metabolism, 41(7), 1563–1578. 10.1177/0271678X20972856

Bashyam, V. M., Erus, G., Doshi, J., Habes, M., Nasralah, I., Truelove-Hill, M., Srinivasan, D., Mamourian, L., Pomponio, R., Fan, Y., Launer, L. J., Masters, C. L., Maruff, P., Zhuo, C., Völzke, H., Johnson, S. C., Fripp, J., Koutsouleris, N., Satterthwaite, T. D., … Davatzikos, C. (2020). MRI signatures of brain age and disease over the lifespan based on a deep brain network and 14 468 individuals worldwide. Brain, 143(7), 2312–2324. 10.1093/brain/awaa160

Basser, P. J., & Pierpaoli, C. (2011). Microstructural and physiological features of tissues elucidated by quantitative-diffusion-tensor MRI. Journal of Magnetic Resonance, 213(2), 560–570. 10.1016/j.jmr.2011.09.022

Bernard, J. A. (2022). Don’t forget the little brain: A framework for incorporating the cerebellum into the understanding of cognitive aging. Neuroscience & Biobehavioral Reviews, 137, 104639. 10.1016/j.neubiorev.2022.104639

Böhle, M., Eitel, F., Weygandt, M., & Ritter, K. (2019). Layer-Wise Relevance Propagation for Explaining Deep Neural Network Decisions in MRI-Based Alzheimer\textquotesingles Disease Classification. Frontiers in Aging Neuroscience, 11. 10.3389/fnagi.2019.00194

Boutinaud, P., Tsuchida, A., Laurent, A., Adonias, F., Hanifehlou, Z., Nozais, V., Verrecchia, V., Lampe, L., Zhang, J., & Zhu, Y.-C. (2021). 3D segmentation of perivascular spaces on T1-weighted 3 Tesla MR images with a convolutional autoencoder and a U-shaped neural network. Frontiers in Neuroinformatics, 15, 641600.

Chollet, F. (2015). *Keras*. https://keras.io/

Cole, J. H., Poudel, R. P. K., Tsagkrasoulis, D., Caan, M. W. A., Steves, C., Spector, T. D., & Montana, G. (2017). Predicting brain age with deep learning from raw imaging data results in a reliable and heritable biomarker. NeuroImage, 163, 115–124. 10.1016/j.neuroimage.2017.07.059

Cox, S. R., Harris, M. A., Ritchie, S. J., Buchanan, C. R., Valdés Hernández, M. C., Corley, J., Taylor, A. M., Madole, J. W., Harris, S. E., Whalley, H. C., McIntosh, A. M., Russ, T. C., Bastin, M. E., Wardlaw, J. M., Deary, I. J., & Tucker-Drob, E. M. (2021). Three major dimensions of human brain cortical ageing in relation to cognitive decline across the eighth decade of life. Molecular Psychiatry, 26(6), 2651–2662. 10.1038/s41380-020-00975-1

Cuadrado-Godia, E., Dwivedi, P., Sharma, S., Ois Santiago, A., Roquer Gonzalez, J., Balcells, M., Laird, J., Turk, M., Suri, H. S., Nicolaides, A., Saba, L., Khanna, N. N., & Suri, J. S. (2018). Cerebral Small Vessel Disease: A Review Focusing on Pathophysiology, Biomarkers, and Machine Learning Strategies. Journal of Stroke, 20(3), 302–320. 10.5853/jos.2017.02922

De Leeuw, F. E., de Groot, J. C., Achten, E., Oudkerk, M., Ramos, L. M. P., Heijboer, R., Hofman, A., Jolles, J., Van Gijn, J., & Breteler, M. M. B. (2001). Prevalence of cerebral white matter lesions in elderly people: A population based magnetic resonance imaging study. The Rotterdam Scan Study. *Journal of Neurology*, Neurosurgery & Psychiatry, 70(1), 9–14.

Desikan, R. S., Ségonne, F., Fischl, B., Quinn, B. T., Dickerson, B. C., Blacker, D., Buckner, R. L., Dale, A. M., Maguire, R. P., Hyman, B. T., Albert, M. S., & Killiany, R. J. (2006). An automated labeling system for subdividing the human cerebral cortex on MRI scans into gyral based regions of interest. NeuroImage, 31(3), 968–980. 10.1016/j.neuroimage.2006.01.021

Dinsdale, N. K., Bluemke, E., Smith, S. M., Arya, Z., Vidaurre, D., Jenkinson, M., & Namburete, A. I. L. (2021). Learning patterns of the ageing brain in MRI using deep convolutional networks. NeuroImage, 224, 117401. 10.1016/j.neuroimage.2020.117401

Duering, M., Biessels, G. J., Brodtmann, A., Chen, C., Cordonnier, C., Leeuw, F.-E. de, Debette, S., Frayne, R., Jouvent, E., Rost, N. S., Telgte, A. ter, Salman, R. A.-S., Backes, W. H., Bae, H.-J., Brown, R., Chabriat, H., Luca, A. D., deCarli, C., Dewenter, A., … Wardlaw, J. M. (2023). Neuroimaging standards for research into small vessel disease—Advances since 2013. The Lancet Neurology, 22(7), 602–618. 10.1016/S1474-4422(23)00131-X

Eberle, O., Buttner, J., Krautli, F., Muller, K.-R., Valleriani, M., & Montavon, G. (2022). Building and Interpreting Deep Similarity Models. IEEE Transactions on Pattern Analysis and Machine Intelligence, 44(3), 1149–1161. 10.1109/TPAMI.2020.3020738

Eitel, F., Soehler, E., Bellmann-Strobl, J., Brandt, A. U., Ruprecht, K., Giess, R. M., Kuchling, J., Asseyer, S., Weygandt, M., Haynes, J.-D., Scheel, M., Paul, F., & Ritter, K. (2019). Uncovering convolutional neural network decisions for diagnosing multiple sclerosis on conventional MRI using layer-wise relevance propagation. NeuroImage: Clinical, 24, 102003. 10.1016/j.nicl.2019.102003

Feng, X., Lipton, Z. C., Yang, J., Small, S. A., & Provenzano, F. A. (2020). Estimating brain age based on a uniform healthy population with deep learning and structural magnetic resonance imaging. Neurobiology of Aging, 91, 15–25. 10.1016/j.neurobiolaging.2020.02.009

Fischl, B., Salat, D. H., Busa, E., Albert, M., Dieterich, M., Haselgrove, C., Van Der Kouwe, A., Killiany, R., Kennedy, D., & Klaveness, S. (2002). Whole brain segmentation: Automated labeling of neuroanatomical structures in the human brain. Neuron, 33(3), 341–355.

Fjell, A. M., & Walhovd, K. B. (2010). Structural brain changes in aging: Courses, causes and cognitive consequences. Reviews in the Neurosciences, 21(3), 187–222.

Fjell, A. M., Westlye, L. T., Amlien, I., Espeseth, T., Reinvang, I., Raz, N., Agartz, I., Salat, D. H., Greve, D. N., Fischl, B., Dale, A. M., & Walhovd, K. B. (2009). High Consistency of Regional Cortical Thinning in Aging across Multiple Samples. Cerebral Cortex, 19(9), 2001–2012. 10.1093/cercor/bhn232

Fjell, A. M., Westlye, L. T., Grydeland, H., Amlien, I., Espeseth, T., Reinvang, I., Raz, N., Holland, D., Dale, A. M., & Walhovd, K. B. (2013). Critical ages in the life course of the adult brain: Nonlinear subcortical aging. Neurobiology of Aging, 34(10), 2239– 2247. 10.1016/j.neurobiolaging.2013.04.006

Francis, F., Ballerini, L., & Wardlaw, J. M. (2019). Perivascular spaces and their associations with risk factors, clinical disorders and neuroimaging features: A systematic review and meta-analysis. International Journal of Stroke, 14(4), 359–371. 10.1177/1747493019830321

Ghassemi, M., Oakden-Rayner, L., & Beam, A. L. (2021). The false hope of current approaches to explainable artificial intelligence in health care. The Lancet Digital Health, 3(11), e745–e750. 10.1016/S2589-7500(21)00208-9

Hofmann, S. M., Beyer, F., Lapuschkin, S., Goltermann, O., Loeffler, M., Müller, K.-R., Villringer, A., Samek, W., & Witte, A. V. (2022). Towards the Interpretability of Deep Learning Models for Multi-modal Neuroimaging: Finding Structural Changes of the Ageing Brain. NeuroImage, 119504. 10.1016/j.neuroimage.2022.119504

Huang, P., Zhu, Z., Zhang, R., Wu, X., Jiaerken, Y., Wang, S., Yu, W., Hong, H., Lian, C., Li, K., Zeng, Q., Luo, X., Xu, X., Yu, X., Yang, Y., & Zhang, M. (2021). Factors Associated With the Dilation of Perivascular Space in Healthy Elderly Subjects. Frontiers in Aging Neuroscience, 13, 624732. 10.3389/fnagi.2021.624732

Jenkinson, M., Beckmann, C. F., Behrens, T. E., Woolrich, M. W., & Smith, S. M. (2012). Fsl. Neuroimage, 62(2), 782–790.

Klapwijk, E. T., Van De Kamp, F., Van Der Meulen, M., Peters, S., & Wierenga, L. M. (2019). Qoala-T: A supervised-learning tool for quality control of FreeSurfer segmented MRI data. NeuroImage, 189, 116–129.

Klein, A., & Tourville, J. (2012). 101 Labeled Brain Images and a Consistent Human Cortical Labeling Protocol. Frontiers in Neuroscience, 6. 10.3389/fnins.2012.00171

Lakens, D. (2022). Improving Your Statistical Inferences (Version v1.0.0) [Computer software]. Zenodo. 10.5281/ZENODO.6409077

Lapuschkin, S., Wäldchen, S., Binder, A., Montavon, G., Samek, W., & Müller, K.-R. (2019). Unmasking Clever Hans predictors and assessing what machines really learn. Nature Communications, 10(1), 1096. 10.1038/s41467-019-08987-4

Levakov, G., Rosenthal, G., Shelef, I., Raviv, T. R., & Avidan, G. (2020). From a deep learning model back to the brain—Identifying regional predictors and their relation to aging. Human Brain Mapping, hbm.25011. 10.1002/hbm.25011

Li, Q., Yang, Y., Reis, C., Tao, T., Li, W., Li, X., & Zhang, J. H. (2018). Cerebral Small Vessel Disease. Cell Transplantation, 27(12), 1711–1722. 10.1177/0963689718795148

Loeffler, M., Engel, C., Ahnert, P., Alfermann, D., Arelin, K., Baber, R., Beutner, F., Binder, H., Brähler, E., Burkhardt, R., Ceglarek, U., Enzenbach, C., Fuchs, M., Glaesmer, H., Girlich, F., Hagendorff, A., Häntzsch, M., Hegerl, U., Henger, S., … Thiery, J. (2015). The LIFE-Adult-Study: Objectives and design of a population-based cohort study with 10,000 deeply phenotyped adults in Germany. BMC Public Health, 15, 691. 10.1186/s12889-015-1983-z

Lohmann, G., Müller, K., Bosch, V., Mentzel, H., Hessler, S., Chen, L., Zysset, S., & von Cramon, D. Y. (2001). LIPSIA--a new software system for the evaluation of functional magnetic resonance images of the human brain. Computerized Medical Imaging and Graphics: The Official Journal of the Computerized Medical Imaging Society, 25(6), 449–457. 10.1016/s0895-6111(01)00008-8

Lynch, K. M., Sepehrband, F., Toga, A. W., & Choupan, J. (2023). Brain perivascular space imaging across the human lifespan. NeuroImage, 271, 120009. 10.1016/j.neuroimage.2023.120009

Maniega, S. M., Valdés Hernández, M. C., Clayden, J. D., Royle, N. A., Murray, C., Morris, Z., Aribisala, B. S., Gow, A. J., Starr, J. M., Bastin, M. E., Deary, I. J., & Wardlaw, J. M. (2015). White matter hyperintensities and normal-appearing white matter integrity in the aging brain. Neurobiology of Aging, 36(2), 909–918. 10.1016/j.neurobiolaging.2014.07.048

Montavon, G., Samek, W., & Müller, K.-R. (2018). Methods for interpreting and understanding deep neural networks. Digital Signal Processing, 73, 1–15. 10.1016/j.dsp.2017.10.011

Mori, S., Wakana, S., Zijl, P. C. M. van, & Nagae-Poetscher, L. M. (2005). MRI Atlas of Human White Matter. Elsevier.

Peng, H., Gong, W., Beckmann, C. F., Vedaldi, A., & Smith, S. M. (2021). Accurate brain age prediction with lightweight deep neural networks. Medical Image Analysis, 68, 101871. 10.1016/j.media.2020.101871

Pierpaoli, C., & Basser, P. J. (1996). Toward a quantitative assessment of diffusion anisotropy. Magnetic Resonance in Medicine, 36(6), 893–906. 10.1002/mrm.1910360612

Ras, G., Xie, N., Van Gerven, M., & Doran, D. (2022). Explainable Deep Learning: A Field Guide for the Uninitiated. Journal of Artificial Intelligence Research, 73, 329–397. 10.1613/jair.1.13200

Ronneberger, O., Fischer, P., & Brox, T. (2015). *U-Net: Convolutional Networks for Biomedical Image Segmentation* (Version 1). arXiv. 10.48550/ARXIV.1505.04597

Samek, W., Montavon, G., Lapuschkin, S., Anders, C. J., & Muller, K.-R. (2021). Explaining Deep Neural Networks and Beyond: A Review of Methods and Applications. Proceedings of the IEEE, 109(3), 247–278. 10.1109/JPROC.2021.3060483

Schilling, K. G., Archer, D., Yeh, F.-C., Rheault, F., Cai, L. Y., Hansen, C., Yang, Q., Ramdass, K., Shafer, A. T., Resnick, S. M., Pechman, K. R., Gifford, K. A., Hohman, T. J., Jefferson, A., Anderson, A. W., Kang, H., & Landman, B. A. (2022). Aging and white matter microstructure and macrostructure: A longitudinal multi-site diffusion MRI study of 1218 participants. Brain Structure and Function, 227(6), 2111–2125. 10.1007/s00429-022-02503-z

Schnake, T., Eberle, O., Lederer, J., Nakajima, S., Schutt, K. T., Muller, K.-R., & Montavon, G. (2022). Higher-Order Explanations of Graph Neural Networks via Relevant Walks. IEEE Transactions on Pattern Analysis and Machine Intelligence, 44(11), 7581–7596. 10.1109/TPAMI.2021.3115452

Swieten, J. C. V., Hout, J. H. W. V. D., Ketel, B. A. V., Hijdra, A., Wokke, J. H. J., & Gijn, J. V. (1991). Periventricular Lesions In The White Matter On Magnetic Resonance Imaging In The Elderly: A Morphometric Correlation With Arteriolosclerosis And Dilated Perivascular Spaces. Brain, 114(2), 761–774. 10.1093/brain/114.2.761

Thomas, A. W., Heekeren, H. R., Müller, K.-R., & Samek, W. (2019). Analyzing neuroimaging data through recurrent deep learning models. Frontiers in Neuroscience, 13, 1321.

Tinauer, C., Damulina, A., Sackl, M., Soellradl, M., Achtibat, R., Dreyer, M., Pahde, F., Lapuschkin, S., Schmidt, R., Ropele, S., Samek, W., & Langkammer, C. (2024). *Explainable concept mappings of MRI: Revealing the mechanisms underlying deep learning-based brain disease classification* (arXiv:2404.10433). arXiv. http://arxiv.org/abs/2404.10433

Tinauer, C., Heber, S., Pirpamer, L., Damulina, A., Schmidt, R., Stollberger, R., Ropele, S., & Langkammer, C. (2022). Interpretable brain disease classification and relevance-guided deep learning. Scientific Reports, 12(1), 20254. 10.1038/s41598-022-24541-7

Tournier, J.-D., Smith, R., Raffelt, D., Tabbara, R., Dhollander, T., Pietsch, M., Christiaens, D., Jeurissen, B., Yeh, C.-H., & Connelly, A. (2019). *MRtrix3*: A fast, flexible and open software framework for medical image processing and visualisation. NeuroImage, 202, 116137. 10.1016/j.neuroimage.2019.116137

Vieira, S., Pinaya, W. H. L., & Mechelli, A. (2017). Using deep learning to investigate the neuroimaging correlates of psychiatric and neurological disorders: Methods and applications. Neuroscience & Biobehavioral Reviews, 74, 58–75. 10.1016/j.neubiorev.2017.01.002

Wardlaw, J. M., Benveniste, H., Nedergaard, M., Zlokovic, B. V., Mestre, H., Lee, H., Doubal, F. N., Brown, R., Ramirez, J., MacIntosh, B. J., Tannenbaum, A., Ballerini, L., Rungta, R. L., Boido, D., Sweeney, M., Montagne, A., Charpak, S., Joutel, A., Smith, K. J., … colleagues from the Fondation Leducq Transatlantic Network of Excellence on the Role of the Perivascular Space in Cerebral Small Vessel Disease. (2020). Perivascular spaces in the brain: Anatomy, physiology and pathology. Nature Reviews Neurology, 16(3), 137–153. 10.1038/s41582-020-0312-z

Wardlaw, J. M., Valdés Hernández, M. C., & Muñoz-Maniega, S. (2015). What are White Matter Hyperintensities Made of? Journal of the American Heart Association, 4(6), e001140. 10.1161/JAHA.114.001140

Zhang, Y.-D., Pan, C., Sun, J., & Tang, C. (2018). Multiple sclerosis identification by convolutional neural network with dropout and parametric ReLU. Journal of Computational Science, 28, 1–10. 10.1016/j.jocs.2018.07.003

